# Assessing actimeters for inclusion in the Healthy Brain Network

**DOI:** 10.1101/183772

**Authors:** Jonathan Clucas, Curt White, Bonhwang Koo, Michael Milham, Arno Klein

## Abstract

**Background:** The Healthy Brain Network is an openly shared pediatric psychiatric biobank with a target of 10,000 participants between the ages of 5 and 21, inclusively In adding ecological actimetry to the Healthy Brain Network, we intend to use appropriate, accurate, reliable tools. Currently a wide range of personal activity trackers are commercially available, providing a wide variety of sensor configurations. For many of these devices, accelerometry provides the basis of measuring both physical activity and sleep with comparable derivative measures.

**Results:** In order to include an ecological biotracker in the Healthy Brain Network protocol, we first evaluated the specifications of a variety of actimeters available for purchase. We then acquired physical instances of 5 of these devices (ActiGraph wGT3X-BT, Empatica Embrace, Empatica E4, GENEActiv Original, and Wavelet Wristband) and wore each of them in our daily lives, annotating our activities and evaluating the reasonableness of the data from each device and the logistical affordances of each device.

**Conclusions:** We decided that the ActiGraph wGT3X-BT is the most appropriate device for inclusion in the Healthy Brain Network. However, none of the devices we evaluated was clearly superior or inferior to the rest; rather, each device seems to have use cases in which that device excels beyond the others.

## Background

The Healthy Brain Network is an openly shared pediatric psychiatric biobank that is an ongoing initiative[1]. In selecting an actimeter for inclusion, we needed to balance data affordances with the practicalities of asking children to wear these devices. Our participants are inclusively 5 to 21 years old, and for inclusion in our biobank we want data as granular and reliable as possible.

A wide range of personal actimeters have been released commercially in recent years. The range of available devices includes research-grade devices, personal trackers, developer kits and toys. “The proliferation of wearable sensing systems has made it possible to continuously record various physiological signals for prolonged periods. … However, it is very difficult to perform a causal analysis on this data in order to find the underlying reasons for differences and trends”[2, p. 268] due to lack of reliable contextual data and the difficulties of temporal harmonization across biosignals. Clocks drift, sampling rates differ and manufacturers make independent decisions regarding factors including signal filtering and units of measurement.

Within a given intended market, prior research indicates “high interdevice reliability”[3, p. 20] within brands and “a high consistency between brands”[4]. Comparing a research-grade ActiGraph and a consumer-grade FitBit, researchers recently found “increasing disagreement between the two devices when the physical activity intensity is increased” in 9-and 10-year-old wearers[5] p. 4]. In a comparison of 9 different actimeters, Rosenberger, Buman, Haskell, McConnell, and Carstensen found good but imperfect results for each device, with each device outperforming the other in terms of reliability at different levels of intensity[6]. These findings encourage considering contextual use in selecting an actimeter.

In their evaluation of a trio of wearable actimeters for a population of middle-aged women, Huberty, Ehlers, Kurka, Ainsworth, and Buman “suggest researchers consider participant acceptability and demand when choosing a wearable sensor”[7, p. 8], i.e. researchers should consider practical factors such as “appearance, comfort, and inconvenience”[7, p. 5] in addition to factors relating to the data a device is able to collect. Keeping both our data requirements and our participants' user experience in mind, we evaluated a practical subset of the current set of available actimeters prior to selecting one for inclusion in the Healthy Brain Network.

## Methods

### Exclusionary criteria

The marketplace of personal actimetry wearable devices is fairly saturated. The marketplace of personal actimetry wearable devices is fairly saturated. With the knowledge that practical constraints including time, money and device availability would limit us to only sample a small proportion of the myriad devices available, we heuristically narrowed our set of trial devices using the following exclusionary criteria.

### Data from devices

Because the goal of the Healthy Brain Network is to collect and share a wide range of relevant data, data access and ability to share our data are primary considerations in the selection of an actimeter. The coarseness of the most granularly available data eliminated some devices (e.g., Fitbit AltaHR[8]) from consideration in favor of devices with greater available information density.

### Wear time affordances

Our use case involves acquiring biosignals from young individuals over the course of a month. Some otherwise well-suited devices require frequent charging (e.g., Apple Watch[9]), which limits the possible comprehensiveness of the acquirable data. In order to ensure the ability to capture both activity data and sleep data, we eliminated devices requiring charging after fewer than 24 hours.

### Target population affordances

Our target population ranges in age from 5 to 21 years inclusively. Considering typical sizes and lifestyles of people in our target age range led us to eliminate many devices from consideration based on child-friendliness factors including device size, water resistance and damage resistance. Some otherwise promising devices (e.g., Samsung Simband) were eliminated from consideration being “not designed to be water-resistant”[10], being limited to adult sizes, and/or having a limited number of devices available (e.g., Totem Health Sensor Devkit[11]).

### Differentiation

Some device manufacturers have several devices with few pairwise differences relevant to our use case (e.g., GENEActiv Original, GENEActiv Action, GENEActiv Sleep and GENEActiv Wireless[12]). In these cases, we chose a single device from the available set to consider.

Using these criteria, we reduced our set of devices under consideration to 5 wrist-worn devices: ActiGraph wGT3X-BT, Empatica Embrace, Empatica E4, GENEActiv Original, and Wavelet Wristband. Though our initial exploration of the set of available devices included devices with other intended body placement, all the devices in our heuristically determined set happen to be wrist-worn devices.

### Evaluation

Three members of our team wore multiple devices simultaneously (see Table 1). While wearing the devices, we manually logged device time on charger, sleep time, physical exercise, emotional activity (e.g., a horror story) and (except for E4, which is not advertised as water resistant) times getting the devices wet (see Appendix).

The primary purposes of our evaluation were to assess the reasonableness of manufacturer claims and to attempt to anticipate our young participants' potential respective experiences with each device.

**Table 1.** Schedule of device wear times for the research team.

| person | wrist | Monday, 4/3 | Tuesday, 4/4 | Wednesday, 4/5 | Thursday, 4/6 | Friday, 4/7 | Saturday, 4/8 | Sunday, 4/9 |
| --- | --- | --- | --- | --- | --- | --- | --- | --- |
| Arno | left |  |  |  |  |  |  |  |
|  | right |  |  | GENEActiv |  | Embrace |  |  |
|  |  |  |  | E4 | Actigraph | E4 |  |  |
|  |  |  |  |  | Wavelet |  |  |  |
| Curt | left | GENEActiv |  |  |  |  |  |  |
|  |  | E4 | Embrace |  |  |  |  |  |
|  | right | Embrace | Actigraph |  |  |  |  |  |
| Jon | left | GENEActiv |  | Actigraph | Embrace |  |  |  |
|  |  | Wavelet | E4 | Wavelet |  |  |  |  |
|  | right | Actigraph | Wavelet | GENEActive |  |  |  |  |
|  |  |  |  | Embrace | E4 |  |  |  |

Three researchers wore multiple actimeters concurrently. When actimeter placements were changed, those changes occurred during early afternoons to allow for consistent overnight wear.

### Wear time affordances

For our intended use case, primary considerations beyond data availability include potential total recording time, potential daily recording time, available sampling rates, and estimated comfort level. We want the devices to record comparable data in as many contexts as possible. To this end, the devices that prioritize battery life over number of different data streams (see Tables 2 and 3) are preferable for our current use case unless one or more of the additional data streams seems likely to greatly increase the value of the combined data.

**Table 2.** Reported battery and memory.

| device | reported battery life | reported memory |  | sampling rate (Hz) |  |  | source of reported values |
| --- | --- | --- | --- | --- | --- | --- | --- |
|  |  | physical | time | min | default | max |  |
| GENEActiv Original | 1 month | 0.5 Gb | 45 days | 10 | 10 | 100 | [13] |
| ActiGraph wGT3X-BT | 25 days | 4 GB | 180 days | 30 | 30 | 100 | [14] |
| Wavelet Wristband | 5 days | [undisclosed] | [undisclosed] | 1 | 10 | 30 | [15] |
| Empatica E4 | 36 hours | [undisclosed] | 60+ hours | 1 | 32 | 64 | [16] |
| Empatica Embrace | 27 hours | 8 Mb | 48 hours | 32 | 32 | 32 | [17][18] |

These values are advertised by the device manufacturers.

**Table 3.** Included sensors.

|  | actigraphy |  | heart rate |  | EDA | light | temperature |
| --- | --- | --- | --- | --- | --- | --- | --- |
|  | accelerometer | gyroscope | PPG | ECG |  |  |  |
| ActiGraph wGT3X-BT | ✓ |  |  | Polar H7 (via Bluetooth) |  | ✓ |  |
| Empatica Embrace | ✓ | ✓ |  |  | ✓ |  | ✓ |
| EmpaticaE4 | ✓ |  | ✓ |  | ✓ |  | ✓ |
| Wavelet Wristband | ✓ | ✓ | ✓ |  |  |  |  |
| GENEActiv Original | ✓ |  |  |  |  | ✓ | ✓ |

Tick marks indicate included sensors; empty cells indicate excluded sensors.

### Sensors

No pair of devices that we considered has an identical array of sensors (see Table 3). Each device includes an accelerometer, though not the same accelerometer (see Figure 1).

**Figure 1.**
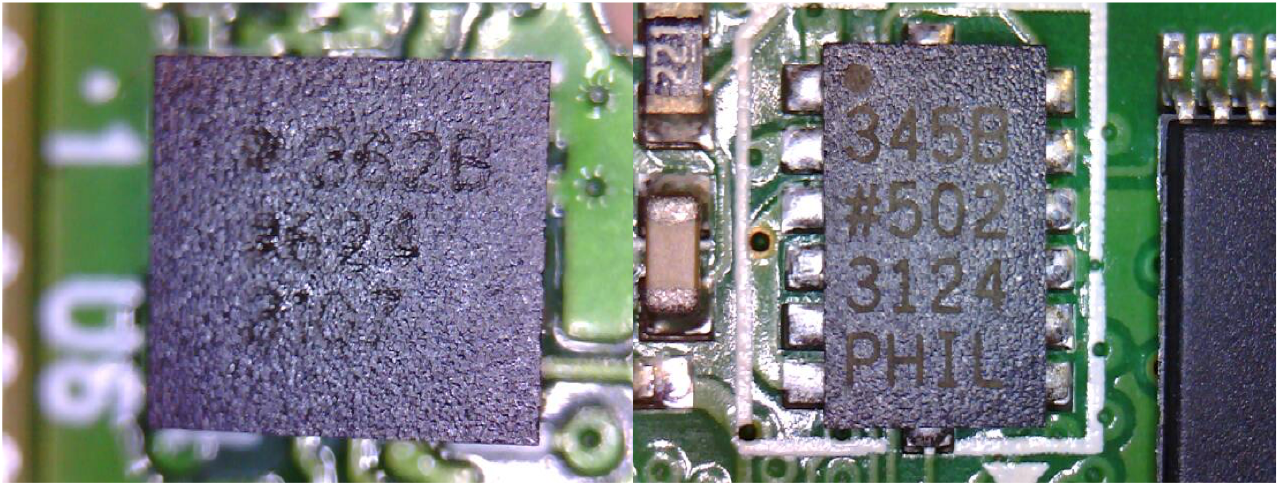
ADXL362B and ADXL345 by Analog Devices. These photographs are of the accelerometers in the ActiGraph wGT3X-BT and one of the GENEActiv Originals that we tested.

Figure 1 depicts ActiGraph vs. GENEActiv in a battle of accelerometers. GENEActiv uses the ADXL345 by Analog Devices[19]. ActiGraph uses the ADXL362, also by Analog Devices[20]. The ADXL362 was released at the same time as the ADXL335, the predecessor of the ADXL345, so ADXL362B is an older chip than ADXL345. The ADXL362 was supposed to be a low power high accuracy chip, while its companion ADXL335 was intended to be ultra low power but not quite as accurate. The ADXL345 improves on the accuracy of the ADXL335 making it comparable in accuracy to the ADXL362 but with the lower power consumption of the ADXL335. The overall difference between the ADXL345 and ADXL362 is negligible.

## Results

Each brand of device has its own specific data format, and each device’s clock has its own drift. Taking these facts into consideration, we attempted to judiciously compare data between these devices.

### Sensors

#### Accelerometry

Research Domain Criteria (RDoC) recommends “longitudinal actigraphy (acrophase, mesor, amplitude)” as an objective measure of the circadian rhythms construct[21, p. 114]. Because all the devices we evaluated include accelerometry as a primary actigraphy sensor, we focused primarily on accelerometry data in our evaluation. Although “a need to standardize accelerometer data reduction” has been noted for decades[22, p. S552], no such standardization has yet been universally adopted. The outputs of each of our test devices seemed reasonable when we acquired data, which was less often than we expected (see Figure 2). Our surprising gaps in data (from times that we made errors in device activation) led us to increase our prioritization of ease of device setup.

**Figure 2.**
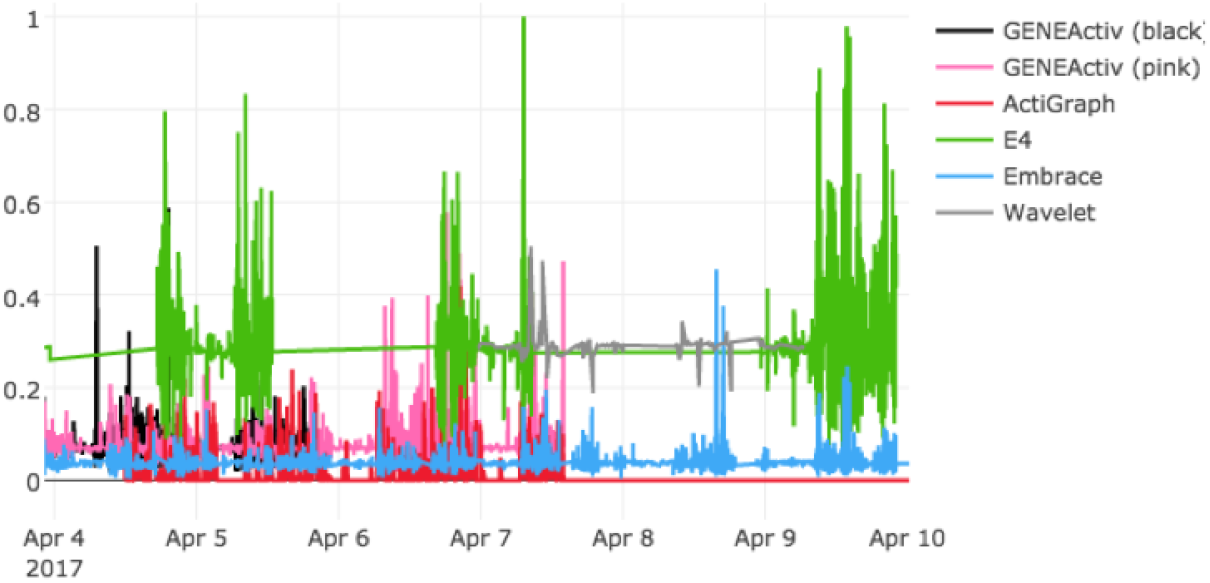
Accelerometry data from all of our devices, April 4-April 10, 2017. The absolute value of each spatial vector (*x, y* and *z*) is normalized against that vector's maximum in the dataset and summed 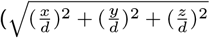 where 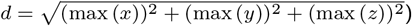 for each time point in an attempt to resolve differences in orientation and magnitude of measurement. Differences in data measurement and/or presentation across devices are still apparent. See Table 1 for a schedule of device use in this timeframe.

Our accelerometry data also illustrated that each brand of device has its own data format, its own clock and its own units of measurement (see Figure 2). In order to compare these data, for each accelerometry stream we calculated from the *x*-axis, *y*-axis and *z*-axis vectors a normalized Euclidian distance from origin vector 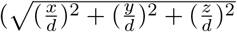 where 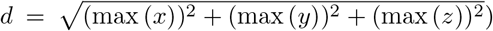, eliminating differences in spatial orientation and minimizing differences in magnitude.

We then conducted a second test (details below) comparing the two most feasible devices.

#### Gyroscopy

Two of the devices we evaluated (Empatica Embrace and Wavelet Wristband) include gyroscopes, but no data were made available to us from those sensors.

#### Photoplethysmography (PPG)

Photoplethysmography uses light reflectance to estimate heart rate. Two of the devices we evaluated (Empatica E4 and Wavelet Wristband) included PPG sensors. However, each of these two devices uses a different set of light colors, and of the two devices, only the Wavelet device provided access to unprocessed PPG data. For these reasons we deemed comparing the PPG data from these devices to each other to be beyond the scope of this evaluation.

#### Electrocardiography (ECG)

Data from an ECG sensor can be recorded with a Bluetooth-connected ActiGraph wGT3X-BT. PPG is also used to measure heart rate, though less directly than ECG (see Figure 3). Our experimental schedule was faulty in that our E4, which provides heart rate measurements from PPG data, and our wGT3X-BT, which provides heart rate measurements from Bluetooth-connected ECG data, never concurrently captured data from the same test subject. Wavelet is capable of providing heart rate measurements as well, but the heart rate measurements from that device are classified as derived rather than raw data.

**Figure 3.**
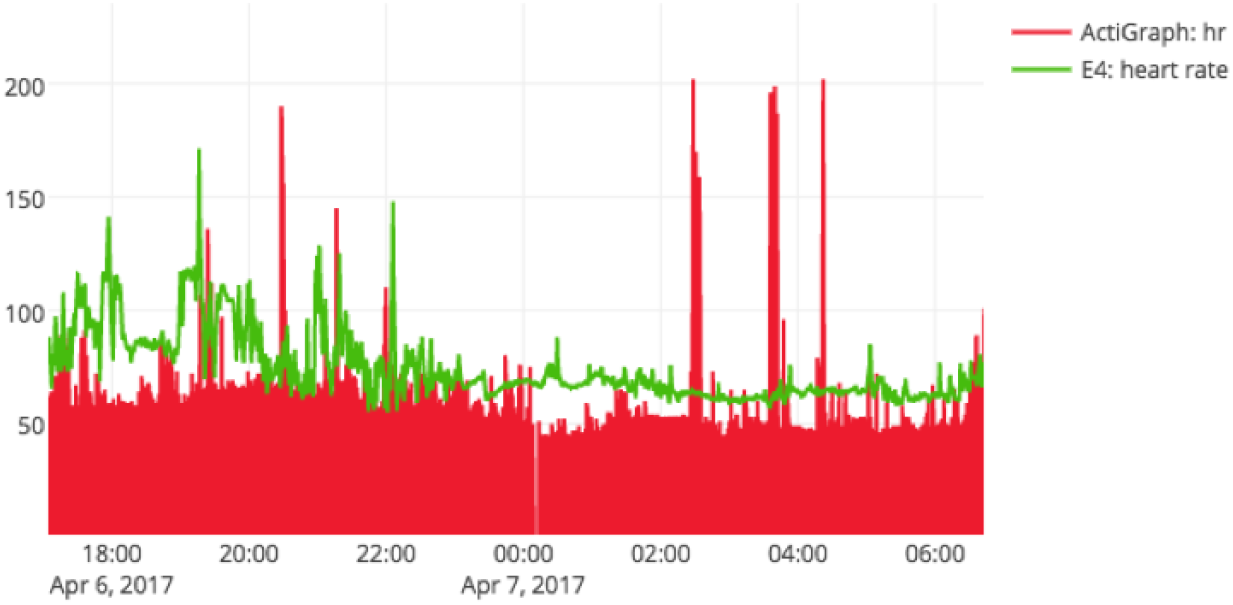
Simultaneous heart rate data from different devices and subjects. The envelopes may be comparable, but the raw data clearly are not.

#### Electrodermal activity (EDA)

EDA sensors are included exclusively on both Empatica devices that we tested. This signal, also known as galvanic skin response (GSR), is advertised as “used to measure sympathetic nervous system arousal and to derive features related to stress, engagement, and excitement”[16]. Skin response measures of EDA have shown both promise and limitations[2]. Our evaluations indicated a coherent EDA signal from each of the Empatica devices (see Figure 4).

**Figure 4.**
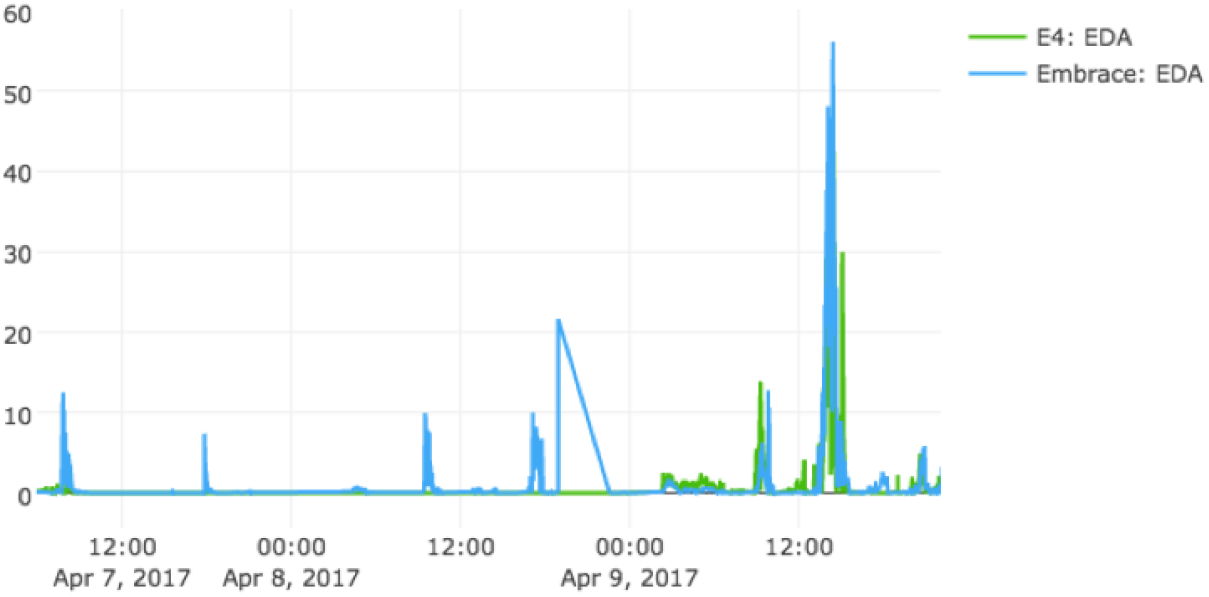
EDA signals from Empatica E4 and Embrace worn concurrently. Although the patterns are not identical, the two Empatica devices appear to be capturing comparable data.

#### Light and Temperature

Neither light nor temperature is a primary feature of interest in our use case. As Figure 5 indicates, our light and temperature sensors captured data during our trials, but in a different format for each brand of device.

**Figure 5.**
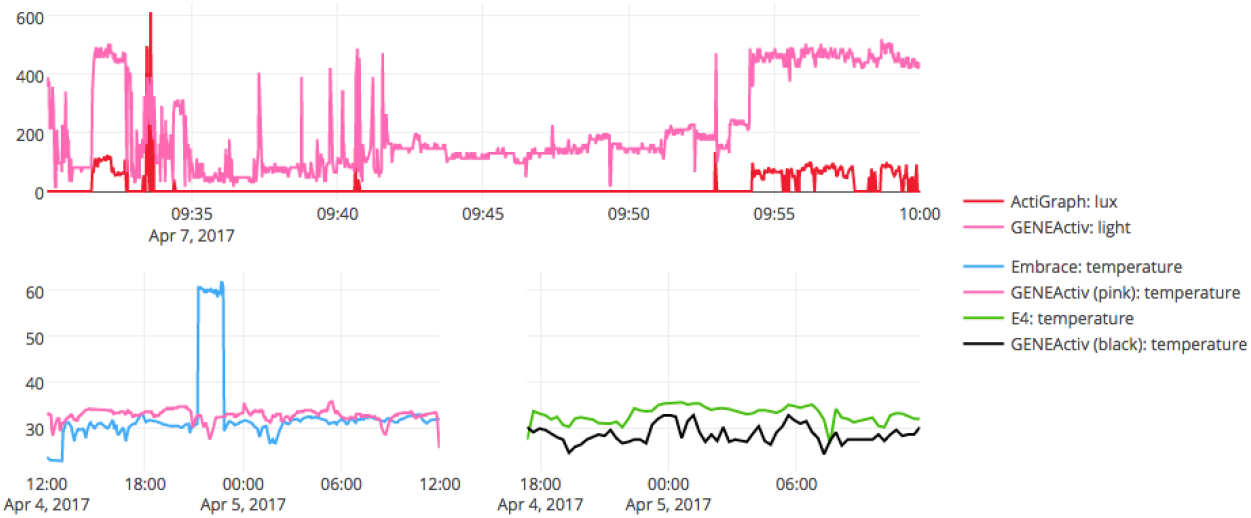
Light readings (top) and temperature readings (bottom). In each example plotted, both devices shown were worn concurrently, side-by-side on the same limb.

The battery life, apparent reliability, ease of set-up and physical design of the ActiGraph wGT3X-BT led us to favor this device over the others for inclusion in the Healthy Brain Network protocol. After making our selection, we performed a follow-up evaluation to assess the comparability of ActiGraph accelerometry data to GENEActiv accelerometry data. GENEActiv also has a long battery life, and we previously used GENEActiv Original in a pilot testing effort for the Healthy Brain Network in which the devices were worn by participants between brain scanning sessions[23]. Past comparisons of ActiGraph actimeters against GENEActiv actimeters found a “strong relation between accelerations measured by the two brands[,] suggest[ing] that habitual activity level and activity patterns assessed by the GENE and GT3X+ may compare well if analyzed appropriately”[24, p. 201]. Although previous research suggests that GENEActiv tends to indicate greater magnitudes of velocity, “it appears that features from the frequency domain are comparable between the two accelerometers[24]”[25, p. 12]. Supporting the temporal comparability of these devices despite differences in magnitude, Hildebrand et al. found “the A[cti]G[raph] and G[ENE]A[ctiv] wrist thresholds in children were different from each other (35 .6 vs. 56.3 mg), despite similar accuracy (AUC = 0.91)”[26, p. 7]. Using previously established cut-points for measuring sedentary behavior in children using wrist-worn actimeters, van Loo et al. found ActiGraph and GENEActiv to both be less than ideal but comparable in classifying sedentary behavior in a sample of 57 5-7 year-old children[27]. Having some GENEActiv Originals and an ActiGraph wGT3X-BT available to test, we conducted a brief follow-up evaluation between these devices in the interest of reliability.

## Follow-up Methods

Having a small collection of GENEActiv devices and a single ActiGraph device to test, we strapped our ActiGraph wGT3X-BT (serial number MOS2D09170617) between two GENEActiv devices (serial numbers 027260 and 028561) on a pair of kali sticks and performed the following actions:

1. held the sticks vertically and churned in a circular path (creating a cylindrical shape) at a roughly consistent velocity;
2. swung the sticks in a horizontal cone at high but roughly consistent velocity;
3. shook the sticks up and down along the axis of the sticks and
4. flung the sticks quickly in a jarring, horizontal V shape.

Each action was 1 minute in duration followed by 1 minute lying motionless on a desk. After the above actions, the 3 trackers were transferred to a researcher’s left (non-dominant) wrist in the same relative placement. Said researcher, who also performed the kali stick actions, performed the following actions while wearing the trackers tightly about his wrist:

1. jumping jacks, about 1 per 1-2 seconds;
2. fast punches, about 1 per second;
3. jogging in place and
4. spinning with arms outstretched.

## Follow-up Results

An initial look at the data supports our suspicion that the devices disagree about the absolute time (see Figure 6). To compare the data streams, we used Holland’s fast normalized cross-correlation script[28] which defines a cross-correlation coefficient *c* of two time series vectors ***x*** and ***y*** with respective lengths *n* and *m* where ***x*** has stride length *m* as 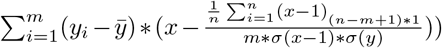. Using this formula, we calculated cross-correlation coefficients of -0.12 and -0.13 between the ActiGraph and each of the GENEActivs and 0.67 between the two GENEActivs.

**Figure 6.**
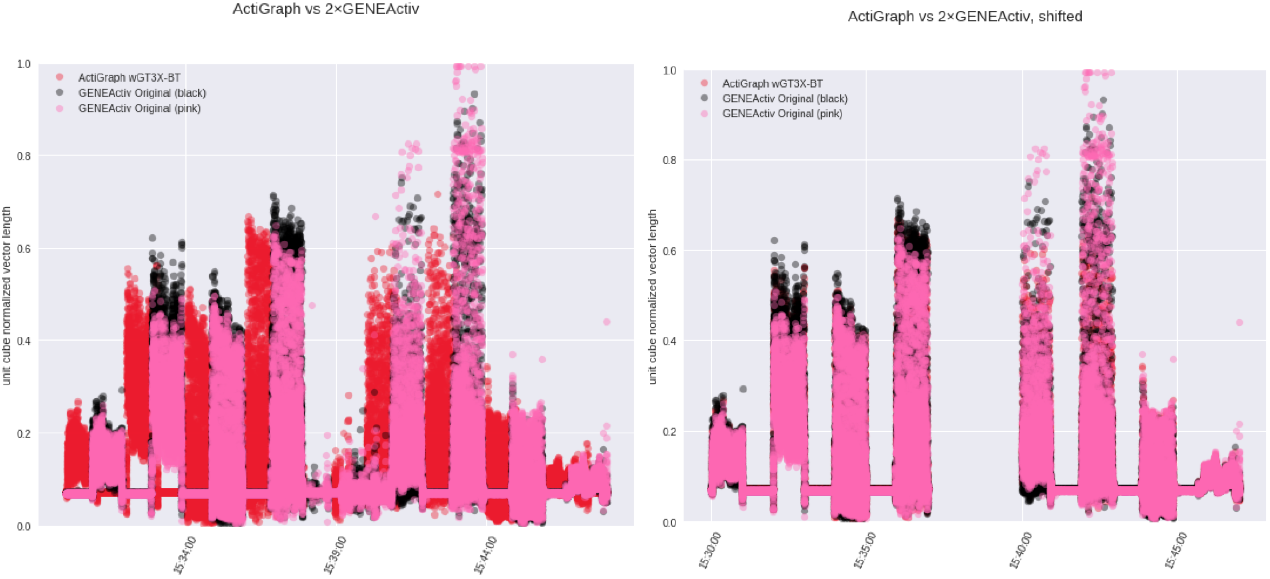
Accelerometry of an Actigraph wGT3X-BT between two GENEActiv Originals. The plot on the left shows the data as timestamped; the plot on the right shows the data with the timestamps shifted.

Computing a translational shift between the ActiGraph and the GENEActivs (a little under 3 seconds) and aligning the three in time (see Figure 6), the resulting normalized cross-correlation coefficients for the ActiGraph and each of the GENEActivs (0.96 and 0.75) are both greater than the normalized cross-correlation between the two GENEActivs (unchanged at 0.67), possibly because the ActiGraph was physically positioned between the two GENEActivs while performing the 8 activities.

Noticeably, the devices indicate acceleration even during the still periods between activities; all 3 devices had mode values around 0.07 normalized vector lengths. We reached out to Activinsights (the manufacturer of GENEActiv), who told us that the “result [we] are seeing is completely normal for the device. This result is due to a small offset in the calibration of the accelerometer”[29], and recommending we use the g.calibrate function in GGIR, including a script to assist in doing so. Applying this script and recalculating the time shift and cross-correlation coefficients, our earlier results are supported with cross-correlation coefficients of 0.64 and 0.97 between the ActiGraph and each of the GENEActivs and 0.66 between the GENEActivs (See Figure 7).

**Figure 7.**
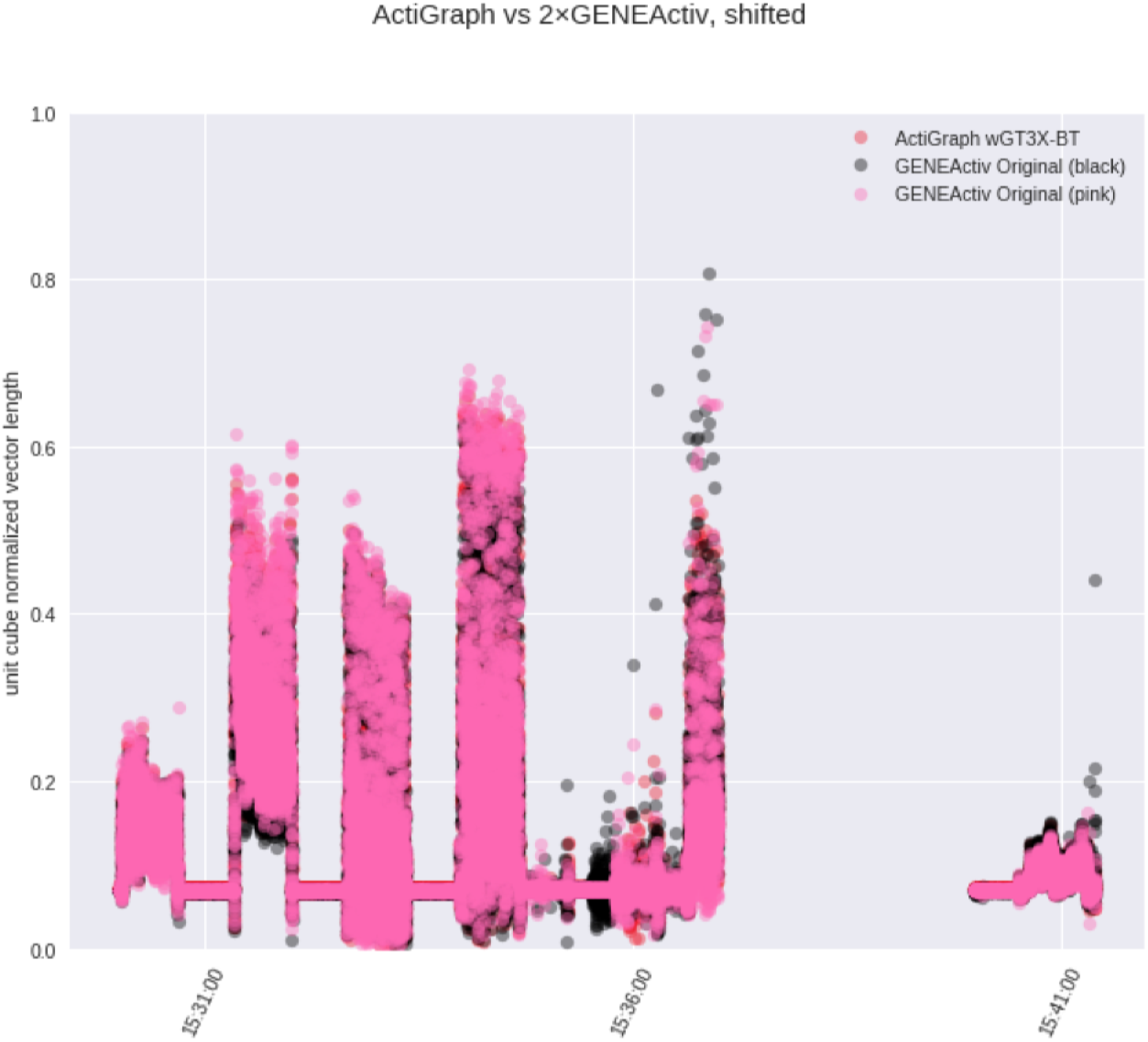
Accelerometry of an Actigraph wGT3X-BT between two GENEActiv Originals. The positive baseline around 0.07 persisted through the GGIR calibration.

## Discussion

Like the data gathered by Rosenberger et al.[6], our data show each device out-performing the others depending on context. While each of the devices showed advantages over the others, the ActiGraph wGT3X-BT afforded the least opportunity for data loss and was for this strength selected as the device to include in the Healthy Brain Network.

### GENEActiv Original

We had a few GENEActiv Original bands on hand from a previous study. The niche for this device and the niche for the ActiGraph wGT3X-BT are largely overlapping. The GENEActiv device includes a temperature sensor in addition to a light sensor and an accelerometer. The software required to initialize the GENEActiv devices and download their data is robust but cumbersome. While these devices would certainly serve the Healthy Brain Network initiative, the GENEActiv ecosystem would more comfortably serve an older (by participant age) and smaller (by number of participants) sample.

### ActiGraph wGT3X-BT

The ActiGraph wGT3X-BT has only an accelerometer, a light sensor and bluetooth for connecting external sensors, a compromise allowing for a long battery life and expandability. The device itself has a small, squat form factor and is a semi-transparent bright red color. After testing all 5 of these devices, our experiences indicated that the wGT3X-BT is the most practical for the Healthy Brain Network’s requirements.

### Wavelet Wristband

Nearly the most comfortable of the devices we evaluated, Wavelet’s Wristband also achieves the median battery life of these 5 actimeters while also providing PPG data by eschewing all other sensors except actigraphy. The Wristband proved to be too comfortable, and with its relatively weak physical strapping mechanism, fell off wrists unnoticed more than once during our in-house tests. Wavelet has been reliably communicative during our evaluation, and the company has updated both their hardware and their software since our evaluation began. For the Healthy Brain Network biobank, we decided the addition of PPG was not worth the risks of device and data loss (from comfortable wristbands and potential forgotten charging sessions). For smaller ecological studies, in terms of time or participants, the Wavelet Wristband would likely be the preferred tracker. Worn together, multiple bands and other same-system devices (i.e., Wavelet Sensor Pods and Wavelet Chest Clips) would provide a rich picture of a participant’s biosignals.

### Empatica E4

While this device provides the richest sensor array of any of the actimeters we evaluated, the E4 also has the shortest battery life. The E4 includes a single onboard button and a single multicolor indicator light which can be used to switch the device on and between live-streaming and recording modes. The potential for confusion of the button-and-light activation coupled with the necessity of asking participants and their families to frequently charge and reactivate the devices renders the E4 impractical for our use-case. The E4 is also the physically largest and most fiscally expensive device that we evaluated. However, with its rich sensor array, the E4 (or upcoming E5) would likely be a preferable biotracker in controlled lab or clinic settings.

### Empatica Embrace

The most comfortable device in our test set, Empatica's Embrace avoids the problems of Wavelet's excess comfort by employing a stretchy band rather than a strap; the Embrace never fell off unnoticed during our tests because the Embrace never fell off at all. Although the Embrace includes a rich sensor array and was also the most stylish of the devices we evaluated, the device requires frequent Bluetooth syncing, and our population of interest is largely too young to be expected to be near enough to a singular paired device every 8 to 9 hours. Additionally, the Embrace’s battery capacity is just over a day. Embrace was engineered to passively detect seizures and is in the process of becoming a general-purpose biotracker; at the time of our evaluation, the raw data access protocols for this device were immature. Once the data access protocols are finalized, the Embrace will likely be an attractive field actimeter for teenagers and adults.

## Conclusions

Rather than selecting an all-purpose device, we selected a device well-suited to the requirements of one particular project. Each of the devices we evaluated, and many that we excluded from our evaluation, have apparent advantages under different requirements. Wavelet released newer, improved versions of both their hardware and their software since we evaluated their Wriststrap, and based on their documentation[15], the new versions reduce or eliminate many of the concerns we raised in our investigation. New devices are being developed by many different organizations while some, such as Jawbone, have ceased production of consumer-grade actimeters in order to focus on medical-grade actimeters[31]; Wavelet itself emerged this way from Amiigo[32]. All of the current actimetry manufacturers continue to release software and hardware updates. In a space as broad and fast-changing as the actimeter market, a common analysis space for each type of sensor would be welcome. Using a common space, care must be taken to specify both how each data set was collected and the ways in which the data were transformed to fit them into the common space.

In addition to adding the ActiGraph wGT3X-BT to the Healthy Brain Network protocol, this evaluation has encouraged us to deeply consider actimeters on a case-by-case basis in future studies. The basic functionality appeared to be adequate in each of the actimeters we evaluated; the differentiating features of these devices can be a boon or a burden depending on the use case.

**List of abbreviations**
ECG: Electrocardiography
EDA: Electrodermal activity
Gb: Gigabit
GB: Gigabyte
Mb: Megabit
PPG: Photoplethysmography
RDoC: Research Domain Criteria

## Declarations

### Ethics approval and consent to participate

No human subjects, human material or human data were involved in this research excepting data from the researchers ourselves.

### Consent for publication

As authors of this study, we consent to publish the data we collected about ourselves.

### Availability of data and material

The datasets generated and/or analysed during the current study are available in the HBN-wearable-analysis repository, https://osf.io/dg869/.

The scripts used for evaluation are available at https://github.eom/ChildMindInstitute/HBN_wearable_evaluation/releases/tag/v1.1.0, including a list of software used.

### Author's contributions

JC participated in all stages of this project, including exploring the universe of available actimeters (1), wearing and evaluating these devices (2), extracting, harmonizing and analyzing the data collected (3), communicating with the device manufacturers, the Healthy Brain Network staff and biosignal experts (4), and drafting and revising this report (5). CW participated in stages 1, 2, 4, 5 and disassembled devices to examine their components when possible (6). Sections of this report were adapted by JC from internal correspondence by CW and AK. BK participated in stages 1, 2 and 5. MM initiated the project and advised, particularly in stages 1, 3 and 5. AK advised and participated in all stages.

## Acknowledgements

We are grateful to Jasmine Escalera^[1]^, Batya Septimus^[1]^ and Lindsay Alexander^[1]^ for logistical support and coordinating the addition of an actimeter into the Healthy Brain Network protocol; to David Scott^[2]^, Rosalind Picard^[3][4]^, Daniel Bender^[4]^ and RJ Kasper^[5]^ for providing us with trial devices and software; and to Charles Sweetland^[6]^ and Sa m Huff^[7]^ for help in interpreting the data we collected.

## Additional File

Additional file 1 — Appendix

A manual log in Microsoft Excel format of device time on charger, sleep time, physical exercise, emotional activity (e.g., a horror story) and (except for E4, which is not advertised as water resistant) times getting the devices wet. This log is not comprehensive and only covers a subset of the time we tested these devices.

## Footnotes

[1] Child Mind Institute, New York, NY, USA

[2] Wavelet Health, Mountain View, CA, USA

[3] MIT Media Lab Affective Computing Group, Cambridge, MA, USA

[4] Empatica, Boston, MA, USA

[5] ActiGraph, Pensacola, FL, USA

[6] Sweetland Solutions Limited, Axmouth, Seaton, Devon, UK

[7] Activinsights Limited, Kimbolton, Cambs, UK

